# Pan-cancer analysis identified inflamed microenvironment associated multi-omics signatures

**DOI:** 10.1101/2020.03.17.996199

**Authors:** Ben Wang, Mengmeng Liu, Zhujie Ran, Xin Li, Jie Li, Yunsheng Ou

## Abstract

**Background:** Immunotherapy has revolutionized cancer therapy. However, responses are not universal. The inflamed tumor microenvironment has been reported to correlate with response in tumor patients. However, how different tumors shape their tumor microenvironment remains a critical unsolved problem. A deeper insight into the molecular characteristics of inflamed tumor microenvironment may be needed.

**Materials and methods:** Here, based on single-cell RNA sequencing technology and TCGA pan-cancer cohort, we investigated multi-omics molecular features of tumor microenvironment phenotypes. Based on single-cell RNA-seq analysis, we classified pan-cancer tumor samples into inflamed or non-inflamed tumor and identified molecular features of these tumors. Analysis of integrating identified gene signatures with a drug-genomic perturbation database identified multiple drugs which may be helpful for converting non-inflamed tumors to inflamed tumors.

**Results:** Our results revealed several inflamed/non-inflamed tumor microenvironments-specific molecular characteristics. For example, inflamed tumors highly expressed miR-650 and lncRNA including MIR155HG and LINC00426, these tumors showed activated cytokines-related signaling pathways. Interestingly, non-inflamed tumors tended to express several genes related to neurogenesis. Multi-omics analysis demonstrated the neuro phenotype transformation may be induced by hypomethylated promoters of these genes and down-regulated miR-650. Drug discovery analysis revealed histone deacetylase inhibitors may be a potential choice for helping favorable tumor microenvironment phenotype transformation and aiding current immunotherapy.

**Conclusion:** Our results provide a comprehensive molecular-level understanding of tumor cell-immune cell interaction and may have profound clinical implications.

## Introduction

The recent clinical successes of immunotherapy represent a turning point in cancer treatment, including immune checkpoint inhibitors, adoptive cell therapy[1, 2]. Clinical trials for anti-PD-1 for patients with melanoma have demonstrated consistent therapeutic responses. Despite these encouraging clinical results, only a fraction of patients can benefit from immunotherapy[3].

Recent studies suggest that the phenotype of tumor microenvironment (TME) is a critical factor influencing the efficacy of immunotherapy. From here, tumor microenvironment can be broadly categorized as being inflamed or non-inflamed tumor microenvironment[4, 5]. Inflamed tumor microenvironment was characterized by rich infiltration of immune cells. These tumors are correlated with the significant tumor regression when treated by immunotherapy[6–10]. However, how different tumor cells shape their TME, thereby determining their response to therapy, remains a critical unsolved problem.

Although some molecular characteristics have been reported in previous researches, a study integrating genome-wide alternations with tumor microenvironment phenotypes would provide deep insights into the mechanisms associated with the formation of inflamed or non-inflamed tumor. To address the question above, we performed comprehensive pan-cancer and multi-omics analysis by comparing inflamed and non-inflamed tumor microenvironment. We identified multiple molecular characteristics associated with tumor immunophenotypes. Analysis of integrating identified gene signatures with drug-genomic perturbation databases uncovers the potentials of multiple drugs on assisting current immunotherapy.

## Results

### Single-cell RNA-seq provides robust genetic signatures of immune cells

To generate deep and robust genetic signatures to dissect tumor microenvironment, we performed single-cell RNA-seq analysis to identify immune cells-specific genetic signatures from a published dataset[11] (Figure.1). Then, these genetic signatures of immune cells were as input for GSVA algorithm[12] to calculate the immune score of tumor samples. This algorithm has been validated in previous reports as an efficient way to evaluate the status of tumor microenvironment[13]. Considering the complexity of tumor microenvironment comprising various types of immune cells, we classified tumor samples into high immune-score(inflamed)/low immune-score(non-inflamed) based on their unsupervised clustering pattern of immune score. Unsupervised clustering algorithm derived from machine learning can group similar entities based on characteristics of data without human intervention. This gives us a chance to yield insights into the complex pattern of tumor microenvironment.

**Figure.1.**
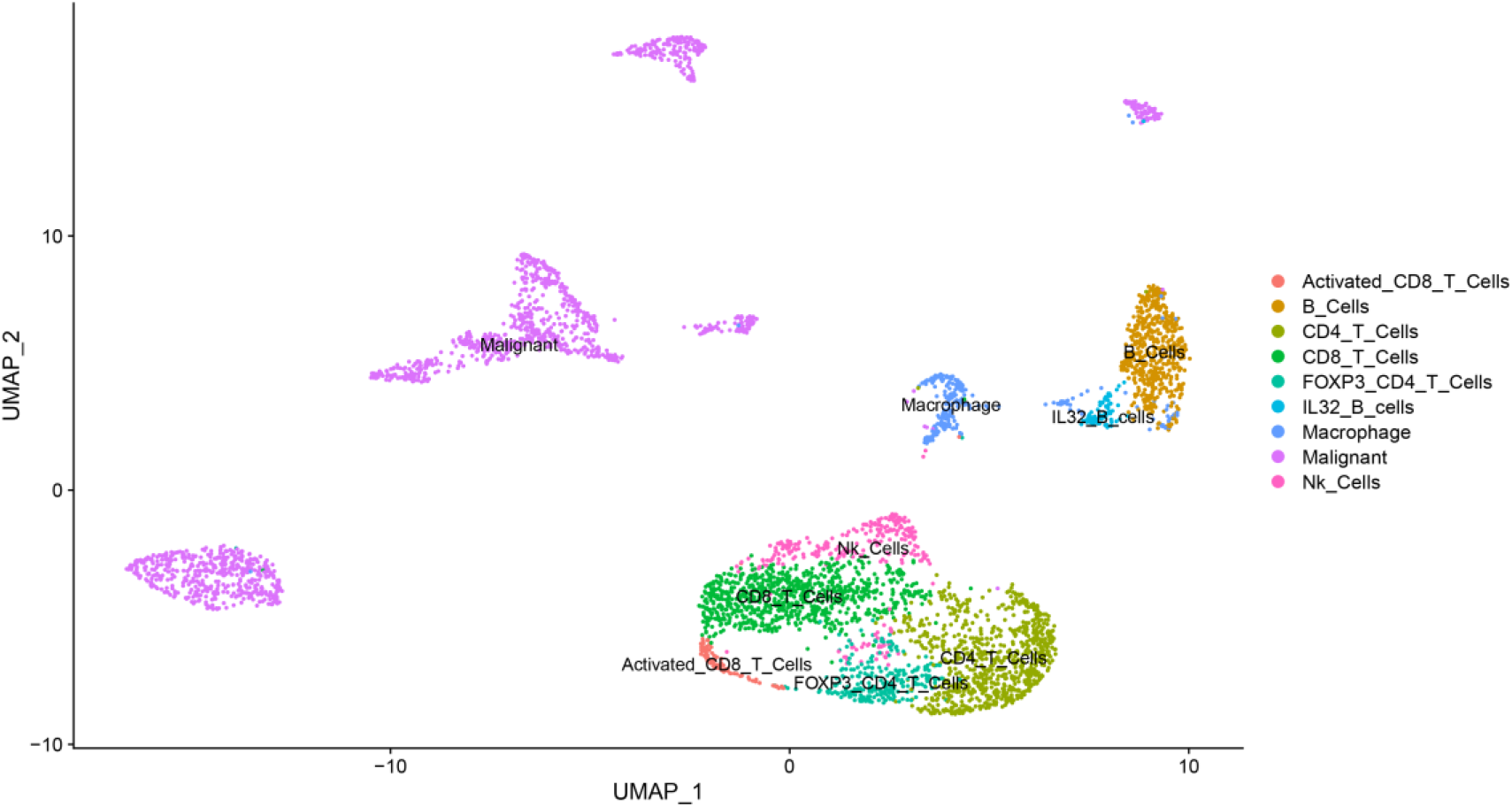
UMAP dimension plot of cells in tumors. Each dot represents a cell colored and labeled by inferred cell types.

### mRNA pattern of inflamed/non-inflamed TME

To validate the robustness of TME classification, *est*imate algorithm was performed to evaluate immune composition of tumor microenvironment[14]. Estimate algorithm infers tumor purity or immune score developed from another independent method. As shown in Figure.2A, high-immune score represents TME phenotype with rich infiltration of immune cells, which is consistent with the results from unsupervised clustering.

**Figure.2.**
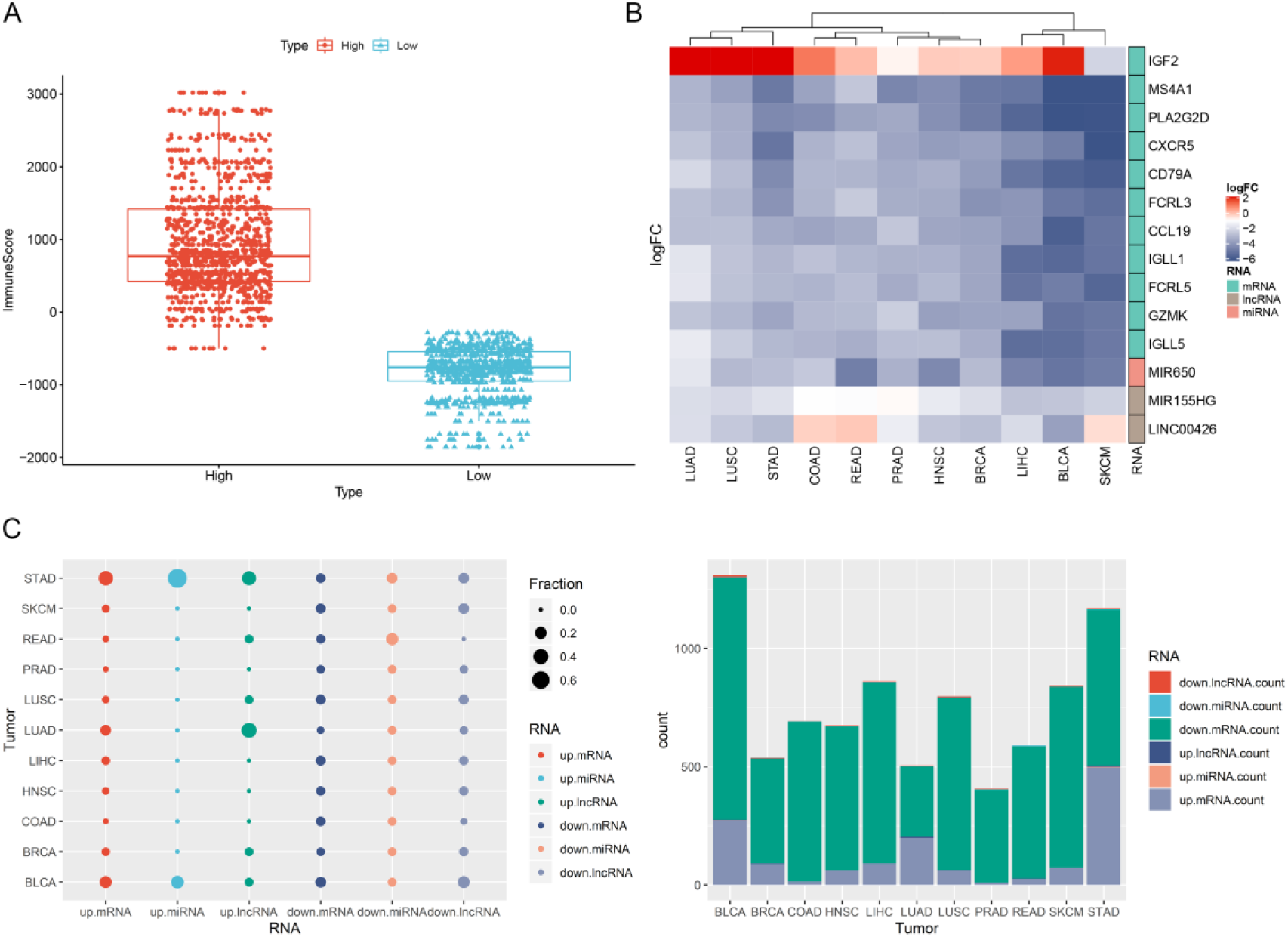
Overview of differently expressed gene between inflamed and non-inflamed TME. A). The high(inflamed)/low-immune score(non-inflamed) TME classification was validated by Estimate algorithm. B). Heatmap for differently expressed mRNA. The cell of heatmap was colored by fold change of genes, red represents genes up-regulated in non-inflamed tumors. C). Overview of molecular signature differences between inflamed and non-inflamed TME

Then, we compared RNA expression between inflamed TME and non-inflamed TME, and significant alternation of RNA expression across different cancer types was observed. The number of differently expressed RNA varied across tumor types (ranging from 406 to 1308, <1000 in BRCA, COAD, HNSC, LIHC, LUAD, LUSC, PRAD, READ, SKCM, >1000 in BLCA, STAD). Among them, down-regulated mRNA (inflamed tumor-specific) represents the most striking signatures that account for major differences between inflamed TME and non-inflamed TME (Figure.2 C).

For example, several genes involved in B cell-associated immune processes were biased at most tumor types (Figure.2B), including Membrane Spanning 4-Domains A1 (MS4A1), Immunoglobulin Lambda Like Polypeptide 1/5 (IGLL1/5), B-Cell Antigen Receptor Complex-Associated Protein Alpha Chain (CD79A), C-X-C Motif Chemokine Receptor 5 (CXCR5), Immune Receptor Translocation-Associated Protein 2 (FCRL5), suggesting inflamed tumor types were also B-cell rich tumor, which has been proven as a key factor determining the sensitivity for immunotherapy[15, 16].

A more systematic functional insight into inflamed/non-inflamed TME demonstrated that genes involved in the neural system, external cellular matrix organization, biological oxidation, IGFBPs-associated pathway were more likely to be upregulated in non-inflamed TME but variate across tumor types (Figure 3A). In contrast, enriched biological processes of inflamed TME-specific mRNA tend to be shared across tumor types: signaling by interleukin and chemokine, PD-1 signaling and co-stimulation by the CD28 family, suggesting a recruiting and inhibitory tumor microenvironment for such inflamed tumor(Figure 3B).

**Figure.3.**
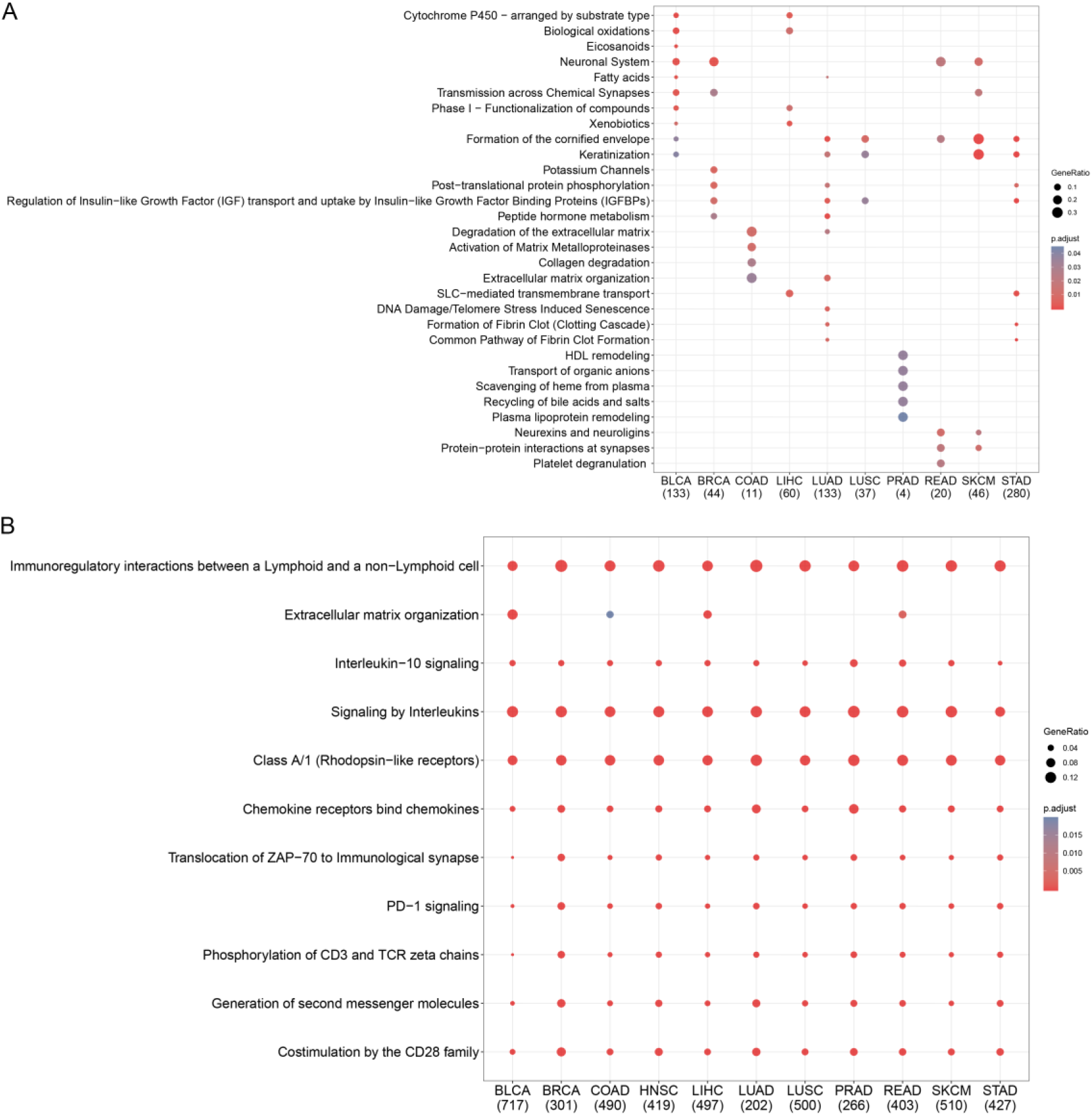
Gene function analysis of inflamed/non-inflamed TME-specific mRNA. A). Enriched biological process for non-inflamed TME-specific expressed gene across tumor types. B). Enriched biological process for inflamed TME-specific expressed gene across tumor types.BLCA, Bladder Urothelial Carcinoma; BRCA, Breast invasive carcinoma; COAD, Colon adenocarcinoma; LIHC, Liver hepatocellular carcinoma; LUAD, Lung adenocarcinoma; LUSC, Lung squamous cell carcinoma; PRAD, Prostate adenocarcinoma; READ, Rectum adenocarcinoma; SKCM, Skin Cutaneous Melanoma; STAD, Stomach adenocarcinoma; HNSC, head and neck squamous cancer;

### non-coding RNA pattern of inflamed and non-inflamed TME

To further investigate the effect of inflamed TME on non-coding RNA expression, we identified miRNAs and lncRNAs that were differently expressed between inflamed TME and non-inflamed TME. Our result demonstrated hsa-miR-650 was up-regulated in inflamed TME in most tumor types (Figure.2B). Its targets were opposing altered in inflamed TME in part of tumors(Figure.4C), including neurogenesis-related genes: CHRNB2, neuronatin (NNAT), neuronal pentraxin 1 (NPTX1), extracellular matrix organization: CAPN6, KRT6A; cellular traffic: KIF1A, SLC34A2(Figure.4A, D). These results revealed that miR-650 may be involved the formation of neuroendocrine phenotype of non-inflamed TME by inhibiting the expression of neurogenesis-related genes[17].

**Figure.4.**
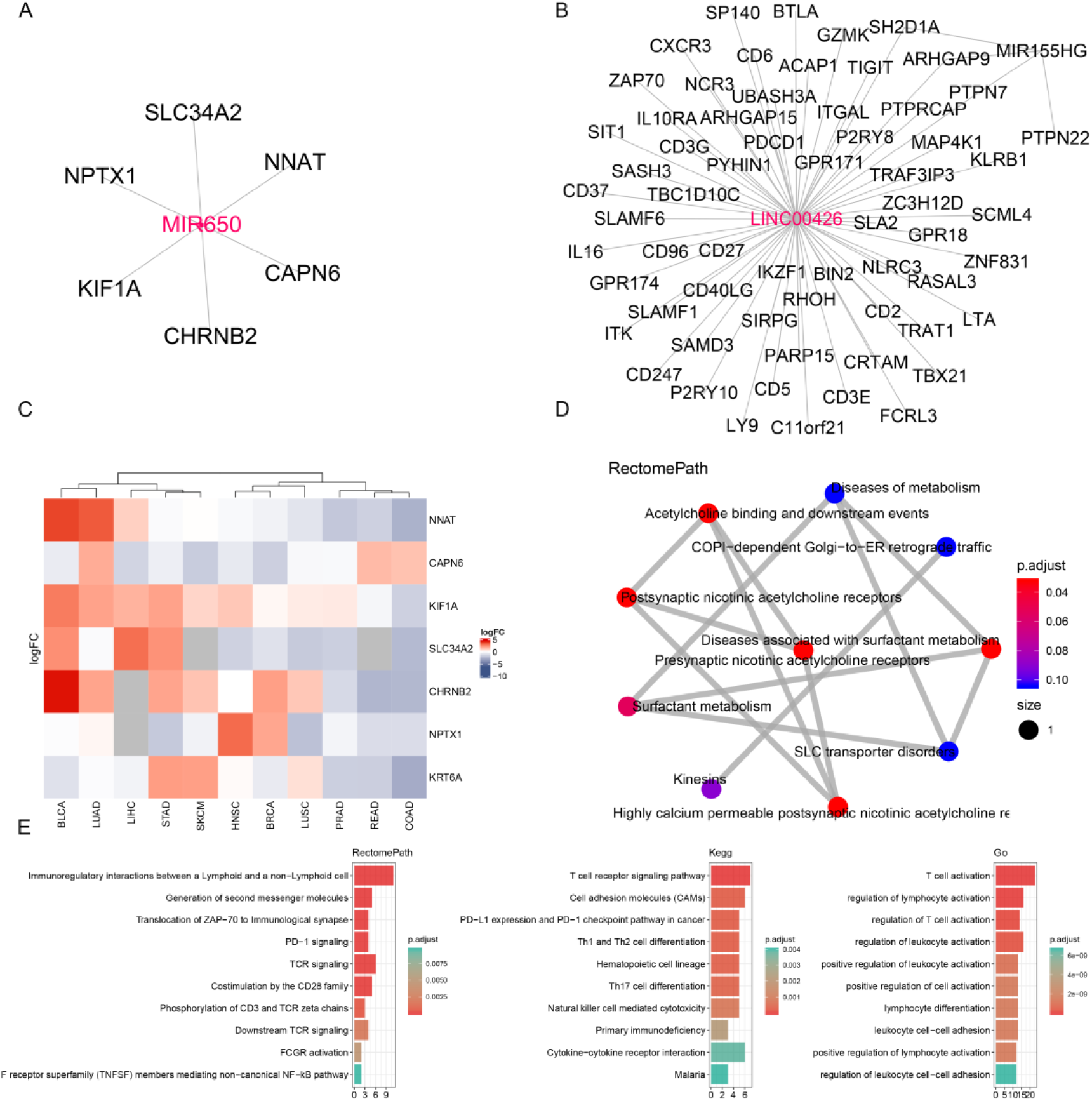
Non-coding RNA pattern of inflamed and non-inflamed TME A). Co-expression network for miR-650 B). Co-expression network for identified lncRNA C). Heatmap for targets of miRNA. The cell of heatmap was colored by fold change of gene. Red represents genes up-regulated in non-inflamed tumors, these genes opposing altered with miR-650 were considered as the targets of miR-650. D). Functional analysis of miRNA targets E). Functional analysis of lncRNA co-expressed genes.

In terms of lncRNA, lncRNAs including MIR155HG and LINC00426 were up-regulated in inflamed TME (Figure.2B). Co-expression analysis revealed that these lncRNAs were associated with multiple immune genes including PDCD1, CXCR3, TIGIT, IL10RA (Figure.4B), which demonstrates that these lncRNAs may play as an immune regulator between lymphoid and non-lymphoid cell (Figure.4E).

### Methylation pattern of inflamed and non-inflamed TME

As well-known, alternation of DNA methylation is an important epigenetic mechanism that results in the dysregulation of RNA. Therefore, we further investigate the alternation of DNA methylation between inflamed and non-inflamed tumors. As shown in the Figure.5A-B, the promoters of several genes involved in oncogenesis biological processes were hypomethylated in non-inflamed tumors, which suggests that the abnormal hypomethylation of these genes may confer the invasive and metastatic ability to non-inflamed tumors. For example, cell-matrix adhesion-related genes: Proto-Oncogene C-Src (SRC), GPI-Anchored Metastasis-Associated Protein Homolog (LYPD3), Mesothelin Like (MSLNL). (supplementary material Table.1)

**Figure.5.**
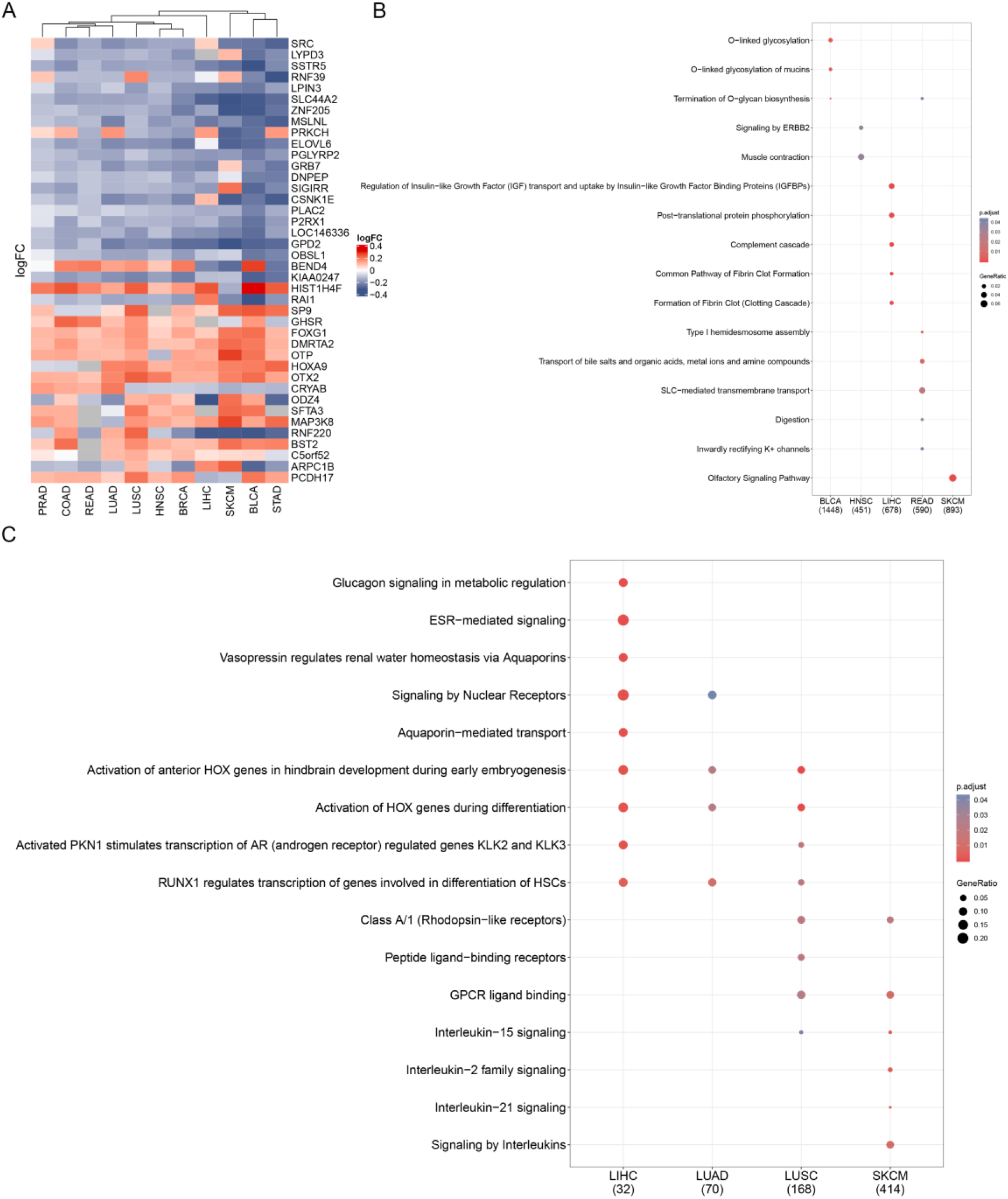
Reactome pathway analysis of inflamed/non-inflamed-specific hypomethylated regions. A). Heatmap for differently methylated probes, grey color represents inflamed tumor-specific hypermethylated probes and non-inflamed tumor-specific hypomethylated probes. Red represent non-inflamed tumor-specific hypermethylated probes and inflamed tumor-specific hypomethylated probes. B). Enriched reactome pathways for non-inflamed tumor-specific hypomethylated gene across TCGA tumor types. C). Enriched reactome pathways for inflamed tumor-specific hypomethylated genes across TCGA tumor types.

Neuroendocrine tumors are dominated by low prevalence of tumor infiltrating lymphocytes with unclear underlying drivers of the non-inflamed phenotype[18, 19]. Consistent with results from mRNA and non-coding RNA, several genes involved in positive regulation of neurogenesis were also hypomethylated in non-inflamed tumors, including Casein Kinase 1 Epsilon (CSNK1E), Protein Kinase C Eta (PRKCH), Obscurin Like Cytoskeletal Adaptor 1 (OBSL1) (supplementary material Table.1), suggesting that hypomethylation of these genes may induce the neuroendocrine phenotype transformation of non-inflamed tumors. Our analysis also revealed promoters of cellular differentiation-related genes including RUNX1 and HOX were hypomethylated across multiple inflamed tumor types (Figure.5C).

These results demonstrated that the hypomethylated of oncogene and neurogenesis-related genes may be an important mechanism involved in the malignant phenotypes transformation of non-inflamed tumors, epigenetic drugs may be promising to shape tumor microenvironment.

### Screening for potential drugs which may convert non-inflamed tumors to inflamed tumors

Despite the understanding of reason why response rate of immunotherapy is unsatisfactory remains limited, it is increasingly clear that combination immunotherapy with targeted therapy is the most feasible way to improve efficacy in a broader patient population[4]. Therefore, it is needed to develop new combination treatment strategies[20].

Here, based on our identified molecular characteristics and a large-scaled drug-genomic perturbation database, we identified multiple drugs that have potentials to convert a non-inflamed into an inflamed tumor (Figure.6A). Two histone deacetylase inhibitors MS275[21] and PHA00816795[22] were identified as the most promising drugs(Figure.6B). More and more evidence has positioned histone deacetylase inhibitors (HDAC) as promising drug candidates by combing it with immunotherapy[23]. However, the effect of histone deacetylase inhibitors on TME remains unclear[24]. Our results suggest that histone deacetylase inhibitors may be a promising way to improve the infiltration ability of immune cells and convert non-inflamed tumors to inflamed one, further clinical trials and experiments are needed to understand the role of histone deacetylase inhibitor.

**Figure.6.**
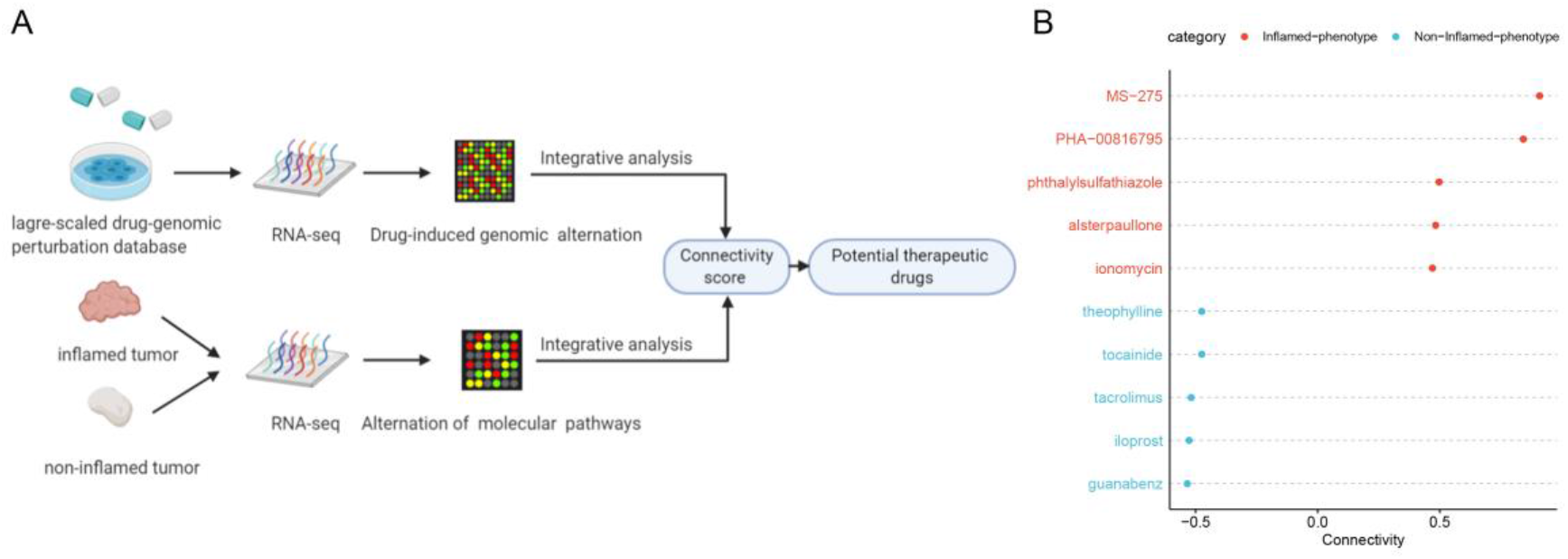
Analysis combining pharmacogenomic perturbation database screens multiple drugs that have the potentials to promote inflamed or non-inflamed immunophenotypic switching. A). The schematic plot shows the workflow of identifying potential drugs that can shape tumor microenvironment. B). The dot plots show potential adjuvant drugs that may contribute to the favorable immunophenotype transformation. Red: Drugs that may induce systematic transcriptomic alternation from non-inflamed TME to inflamed TME. Blue: opposite drugs.

### Identified non-coding RNA and immunophenotypes correlate with tumor patients-prognosis

To explore the prognostic significance of identified non-coding RNA and immune subtypes, univariate/multivariate cox-test and Kaplan-Meier analysis were performed. Kaplan-Meier analysis revealed that these non-coding RNAs were positively associated with better OS in BRCA, LUAD, HNSC, SKCM(Figure.7A). Univariate/Multivariate cox-test also validated the prognostic independence and significance of identified non-coding RNA. (**lnc00426** in BRCA (Univariate: HR=0.7, 95%CI: 0.51-0.95, p=0.024, Multivariate: HR=0.72, 95%CI: 0.53-0.98, p=0.035), HNSC (Univariate: HR=0.7, 95%CI: 0.51-0.95, p=0.024, Multivariate: HR=0.72, 95%CI: 0.53-0.98, p=0.035), LUAD (Univariate: HR=0.74, 95%CI: 0.56-0.98, p=0.034, Multivariate: HR=0.73, 95%CI: 0.55-0.96, p=0.027); **miR155HG** in HNSC (Univariate: HR=0.77, 95%CI: 0.64-0.93, p=0.0068, Multivariate: HR=0.48, 95%CI: 0.32-0.71, p=0.00027), SKCM (Univariate: HR=0.75, 95%CI: 0.67-0.85, p<0.001, Multivariate: HR=0.84, 95%CI: 0.72-0.98, p=0.025); **miR650** in BRCA (Univariate: HR=0.92, 95%CI: 0.87-0.97, p=0.0044, Multivariate: HR=0.72, 95%CI: 0.53-0.98, p=0.035), HNSC (Univariate: HR=0.92, 95%CI: 0.88-0.96, p<0.001, Multivariate: HR=0.48, 95%CI: 0.32-0.71, p<0.001), LUAD (Univariate: HR=0.93, 95%CI: 0.87-1, p=0.043, Multivariate: HR=0.73, 95%CI: 0.55-0.96, p=0.027), SKCM (Univariate: HR=0.93, 95%CI: 0.89-0.98, p=0.003, Multivariate: HR=0.84, 95%CI: 0.72-0.98, p=0.025);)

**Figure.7.**
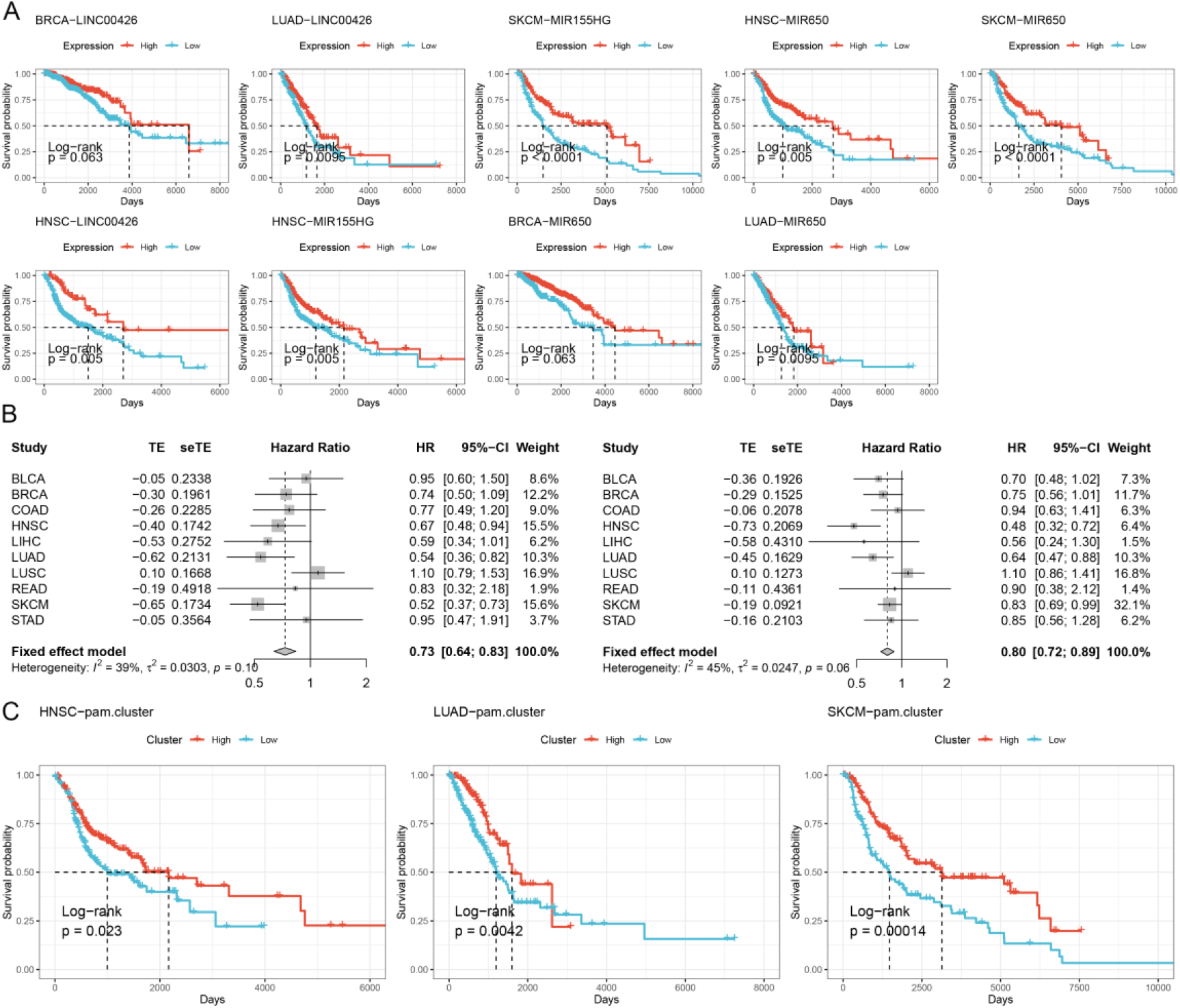
Prognostic role of immunophenotypes and non-coding RNA. A). Kaplan-Meier plots show overall survival rate for the non-coding RNA. The P-value was calculated using log-rank test B). Forest plots show hazard ratio for identified immunophenotypes. the hazard ratio of left panel is calculated by single variable cox test. The right one is from multiple variable cox-test. C). Kaplan-Meier plots show overall survival rate for identified immunophenotypes. The P-value was calculated using log-rank test

We further investigated the prognostic role of immunophenotypes across tumor types. As shown in Figure. 7B, the pool hazard ratio of inflamed TME is 0.73 (Univariate cox test 95%CI 0.64-0.83) or 0.80 (Multivariate cox test,95%CI 0.72-0.89) with not significance heterogeneity (*I*^*2*^=39%, P value= 0.10 in Univariate cox test, *I*^*2*^=45%, P value =0.06 in Multivariate cox test). Kaplan Meier analysis also suggested that inflamed phenotype was associated with better OS (Figure. 7C).

## Discussion

Our study comprehensively revealed many molecular differences and pathways alternation between inflamed and non-inflamed tumor microenvironment across multiple tumor types, such as the chemokine, neurogenesis-related genes.

Compared to inflamed tumors, non-inflamed tumors expressed more neurogenesis-associated genes. The high expression of these genes may confer neuroendocrine-like characteristics to non-inflamed tumors. Neuroendocrine tumors are dominated by low prevalence of tumor infiltrating-lymphocytes with unclear underlying drivers of the non-inflamed phenotype[18, 19]. Our results may provide some mechanistic insights for the transdifferentiation from non-neuroendocrine (inflamed) to neuroendocrine-like (non-inflamed) tumor. For example, miR-650, which is down regulated in non-inflamed tumors, may be involved in the formation of neuroendocrine-like tumor by enhance the expression of its target genes (neurogenesis-associated). Epigenetic regulation may be another potential mechanism involved in the maintenance of neuroendocrine-like phenotype in non-inflamed tumor. Our results demonstrated the promoters of multiple neurogenesis-associated genes were hypomethylated in non-inflamed tumors, which suggests epigenetic regulation may be an important mechanism maintaining the “immune desert” status of non-inflamed tumor and epigenetic drugs such as histone deacetylase inhibitor[25], DNA methylation inhibitors[26, 27], may be a promising choice for converting the immunophenotype of tumor.

Our studies also revealed that the lncRNA may play a general regulator in the inflamed tumor microenvironment, the mRNA expression of MIR155HG and LINC00426 were pan-cancer altered, and the co-expressed genes of MIR155HG and LINC00426 was involved in multiple immune associated biological processes, including immune cell differentiation and exhausted/activated genetic program of T cells, indicating that these lncRNAs might assist the maintenance of inflamed tumor microenvironment across tumor types, especially MIR155HG, which is inflamed TME-specific gene in all of analyzed tumor types. Further researches will elucidate the function of these lncRNA in inflamed TME formation.

To overcome the immunotherapeutic resistance, multiple combination immunotherapy strategies were proposed to improve current response rate of immunotherapy. Here, we identified multiple drugs which may promote favorable TME transformation and provide some clues to better cooperate chemotherapy with current immunotherapy. These drugs-discovery results suggested HDAC inhibitors may be a promising drug to shape TME, which induced inflamed genomic alternation. Further clinical trials will be performed to validate our finding.

This study has some limitations. First, validation in other tumor types or a large cohort is warranted in further study. Second, TCGA doesn’t provide direct status of the tumor microenvironment. Therefore, we had to indirectly infer the relative score of immune composition as described in previous studies[13] and validated by another algorithm[14]. Third, our study provided a comprehensive catalogue of molecular alternations, but we couldn’t further investigate the role of identified molecule for the lack of funding support and experimental environment. Therefore, further studies are necessary to elucidate the detailed role of these molecular characteristics.

In conclusion, our study identified multiple molecular differences between inflamed and non-inflamed TME, these results provide comprehensive insights into inflamed tumor microenvironment-related molecular mechanism and have profound clinical implications: these results may help to optimize current combination immunotherapy to benefit more tumor patients. Nevertheless, our study calls attention to the need to include tumor microenvironment status in future clinical trials.

## Materials and methods

### Multi-omics data for TCGA samples

RNA sequencing data, including mRNA expression, miRNA and lncRNA was downloaded from GEO database with accession number GSE62944[28]. Updated clinical data and DNA methylation data were downloaded from *TCGAbiolinks* [29–31]. The raw count data of RNA sequencing was normalized and quantitated by the edgeR package[32].

### Identifying genetic signatures of immune cell from a Single cell RNA sequencing data

Raw single-cell RNA sequencing data were downloaded from GEO database with accession number GSE72056[11]. This dataset contains 4645 single cells isolated from 19 patients and profiling immune and malignant cells within tumor microenvironment. We applied Seurat packages to normalize the data and identify differentially expressed genes. UMAP was used for dimension reduction. Cells were represented as a dot in a two-dimensional UMAP plane. The annotation of cells was annotated based on raw annotation[11]. Genetic signatures of immune cells were selected as following criteria: 1. Expressed low in malignant cells with proportion of expression <0.1 2. Expressed high in immune cells with proportion of expression >0.5 3. Adjusted p-value <0.05 and log fold change >0

### Classification of TME phenotypes across different tumor types

To investigate the molecular characteristics of inflamed or non-inflamed immunophenotypes, immune cell signatures identified in above single cell RNA sequencing data were used as input for GSVA algorithm[12] to calculate immune score for each immune cell type. GSVA algorithm has been proven as an efficient way to reveal characteristics of tumor microenvironment[13]. Then, tumor samples were classified into low-immune score(non-inflamed), intermediate immune score and high immune score(inflamed) TME groups based on unsupervised clustering pattern of immune score by optCluster [33]. To avoid confounding factors from potential mixture, we excluded samples from the intermediate-immune score from further analysis.

### Identification of molecular differences between inflamed and non-inflamed tumor

Then, we compared the molecular data between these two groups to identify molecular differences. The statistically significant criteria for each molecular characteristic are as follows: mRNA, miRNA and lncRNA: |log FC| >1.5 and *P* value < 0.05. Dysregulated at least four tumor types. We next calculated Pearson correlation coefficient between identified non-coding RNA and coding RNA. Potential coding RNA targets was selected based on criteria as follows: lncRNA: absolute value of correlation coefficient >0.78, P value < 0.05 and dysregulated at least three tumor types. miRNA: validated targets of miRNA[34] (from fourteen miRNA-mRNA interaction database) and opposite alternation of corresponding miRNA. Functional enrichment analysis of potential targets was performed to contribute the mechanical understanding of identified non-coding RNA[35].

In methylation analysis, only CpG probes mapped at promoter regions (e.g. TSS1500, TSS200,5’UTR, 1stexon) were included in this analysis. Differently expressed regions were identified by champ[36] pipeline based on criteria as follows: 1. adjusted P value <0.001 2. The absolute value of beta-value differences >0.1 3. 0.3 was considered as threshold for clearly methylated region.

### Identification of potential drugs that converting a non-immune into an inflamed tumor

Combination immunotherapy is considered as the most efficient way to aid current immunotherapy. Here, transcriptomic differences between inflamed and non-inflamed tumor were used as an input to calculate the connectivity score between transcriptomic differences and drugs-induced genomic alteration. Drugs-induced systematic genomic alteration was from CMap database[37]. The data download and connectivity score calculation were performed with PharmacoGx [38]

### Statistical analysis

The Chi-square test is used for statistical tests of categorical variables. The Kaplan-Meier analysis and log-rank test were used to test survival differences between identified groups. The univariate Cox proportional hazard ratio was used to calculate the hazard ratio of the interested factor. The multivariate Cox proportional hazard ratio was used to test the independence of variate. All statistical analyses were performed by R 3.61.

## Contributions

All authors were involved in the manuscript preparation. In addition, BW: performed the experiments, BW and ML.: Assisted to perform experiments, BW, ZR: designed the summary cartoon, BW, XL, JL: participated the study design, BW and YO: design the study and wrote up the manuscript.

## Acknowledgements

Thanks for everyone who is trying to push the boundary of science.

## Data availability

The data sets generated during and analyzed during the current study are available from the corresponding author upon reasonable request.

